# Enabling cross-study analysis of RNA-Sequencing data

**DOI:** 10.1101/110734

**Authors:** Qingguo Wang, Joshua Armenia, Chao Zhang, Alexander V. Penson, Ed Reznik, Liguo Zhang, Thais Minet, Angelica Ochoa, Benjamin E. Gross, Christine A. Iacobuzio-Donahue, Doron Betel, Barry S. Taylor, Jianjiong Gao, Nikolaus Schultz

**Author notes:** Corresponding author: Nikolaus Schultz.

## Abstract

Driven by the recent advances of next generation sequencing (NGS) technologies and an urgent need to decode complex human diseases, a multitude of large-scale studies were conducted recently that have resulted in an unprecedented volume of whole transcriptome sequencing (RNA-seq) data. While these data offer new opportunities to identify the mechanisms underlying disease, the comparison of data from different sources poses a great challenge, due to differences in sample and data processing. Here, we present a pipeline that processes and unifies RNA-seq data from different studies, which includes uniform realignment and gene expression quantification as well as batch effect removal. We find that uniform alignment and quantification is not sufficient when combining RNA-seq data from different sources and that the removal of other batch effects is essential to facilitate data comparison. We have processed data from the Genotype Tissue Expression project (GTEx) and The Cancer Genome Atlas (TCGA) and have successfully corrected for study-specific biases, enabling comparative analysis across studies. The normalized data are available for download via GitHub (at https://github.com/mskcc/RNAseqDB).

## Introduction

RNA sequencing (RNA-seq) is an important tool for understanding the genetic mechanisms underlying human diseases. A multitude of large-scale studies have recently generated an unprecedented volume of RNA-seq data. For example, The Cancer Genome Atlas (TCGA) has quantified gene expression levels in >8000 samples from >30 cancer types. On a similar scale, the Genotype Tissue Expression (GTEx) project [1], [2], has catalogued gene expression in >9,000 samples across 53 tissues from 544 healthy individuals.

These resources offer a unique opportunity to gain better insight into complex human diseases. However, the integrative analysis of these data across studies poses great challenges, due to differences in sample handling and processing, such as sequencing platform and chemistry, personnel, details in the analysis pipeline, etc. For example, the RNA-seq expression levels of the majority of genes quantified are in the range of 4-10 (log2 of normalized_count) for TCGA, and 0-4 (log2 of RPKM) for GTEx (**Fig. S1A**), a consequence of the use of different analysis pipelines. This makes gene expression levels from the two projects not directly comparable.

To facilitate research on abnormal gene expression in human diseases, a variety of databases and pipelines have been developed to combine RNA-seq from different studies [3][4][5][6][7][8][9][20]. However, these databases or pipelines either directly incorporated expression data from the literature, retaining unwanted batch effects in the data [7][8], or only combined and reanalyzed samples from smaller studies, hence, not taking advantage of the power provided by the recent large data sets [3][4][5][6][20]. A recently published pipeline, Toil [10], attempts to unify RNA-seq data from different sources by uniformly processing raw sequencing reads. However, Toil does not remove batch effects that are introduced by sources other than the differences in read alignment and quantification. To take full advantage of the large volume of available RNA-seq data, an integrative RNA-seq resource is still urgently required.

Here, we present a pipeline for processing and unifying RNA-seq data from different studies. By unifying data from GTEx and TCGA, we provide reference expression levels across the human body for comparison with the expression levels found in human cancer. Our method removes batch effects by uniformly reprocessing RNA-seq data. Specifically, we used raw sequencing reads of the RNA-seq samples downloaded from GTEx and TCGA, realigned them, re-quantified gene expression, and then removed biases specific to each study.

## Results

Our analysis pipeline included realignment of raw reads, removal of degraded samples, expression quantification, and batch effect processing (**Fig. 1**, see **Methods**).

**Figure 1.**
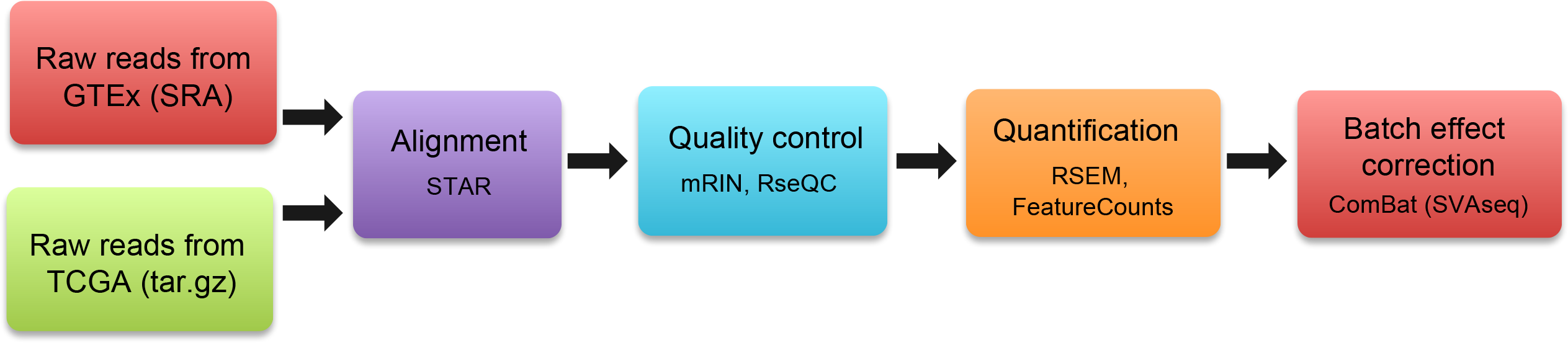
Uniform processing of RNA-seq data from GTEx and TCGA.

To allow proper batch bias correction, we processed only samples from tissues that were studied by both GTEx and TCGA (**Table 1**). Tissues with no or insufficient numbers of normal samples available in TCGA (e.g., sarcoma, ovarian cancer, melanoma) were not processed (**Table S1**).

**Table 1.** GTEx and TCGA RNA-seq samples processed by our pipeline. Only paired-end RNA-seq samples were included.

| GTEx tissue / TCGA cancer type | GTEx | TCGA normal | TCGA tumor | Total |
| --- | --- | --- | --- | --- |
| bladder / blca | 11 | 19 | 411 | 441 |
| breast / brca | 218 | 114 | 1112 | 1444 |
| cervix / cesc | 11 | 3 | 304 | 318 |
| uterus / ucec | 90 | 24 | 180 | 294 |
| uterus / ucs |  | 0 | 57 | 57 |
| colon-sigmoid / read | 173 | 10 | 94 | 277 |
| colon-transverse / coad | 203 | 41 | 295 | 539 |
| liver / lihc | 136 | 50 | 371 | 557 |
| Salivary Gland / hnscc | 70 | 44 | 520 | 634 |
| esophageal / esca | 790 | 11 | 185 | 986 |
| prostate / prad | 119 | 52 | 497 | 668 |
| stomach / stad | 204 | 35 | 415 | 654 |
| thyroid / thca | 355 | 59 | 505 | 919 |
| lung / luad | 374 | 59 | 528 | 961 |
| lung / lusc |  | 51 | 504 | 555 |
| kidney cortex / kirc | 36 | 72 | 541 | 649 |
| kidney cortex / kirp |  | 32 | 290 | 322 |
| kidney cortex / kich |  | 25 | 66 | 91 |
| **Total** | **2790** | **701** | **6875** | **10366** |

We downloaded and processed raw paired-end RNA-seq data from 10,366 samples, including 2,790 from GTEx and 7,576 from the TCGA project (**Table 1**). 831 samples (8%) exhibited 5’ degradation (as described previously [11]) and were excluded from further analysis. We also discarded samples with low alignment rates and samples not used in the final GTEx study, resulting in a total of 9109 (89%) high-quality samples for further analysis.

To correct for batch biases, we first created a sample-gene matrix for each tissue-tumor pair by merging gene expression levels of the corresponding GTEx and TCGA samples. Regardless of the actual batch that a sample belonged to in an RNA-seq experiment, we treated all GTEx samples as one batch and TCGA samples as another. Then, we ran ComBat in the R package SVAseq [12] to correct for non-biological variation accounting for unwanted differences between GTEx and TCGA samples of a particular tissue type (see **Methods**).

To examine how well our pipeline was able to correct study-specific batch effects, we systematically compared the effects of uniform realignment, expression quantification, and batch effect correction for three tissues: bladder, prostate and thyroid. When using expression levels reported by the TCGA and GTEx projects, even after applying upper-quartile normalization to bring expression levels into comparable ranges (**Fig. S1B**), samples from the same study were more similar to each other than samples from the same tissue, as shown by PCA analysis (**Fig. 2A**). This result indicates the necessity to uniformly reprocess RNA-seq samples.

**Figure 2.**
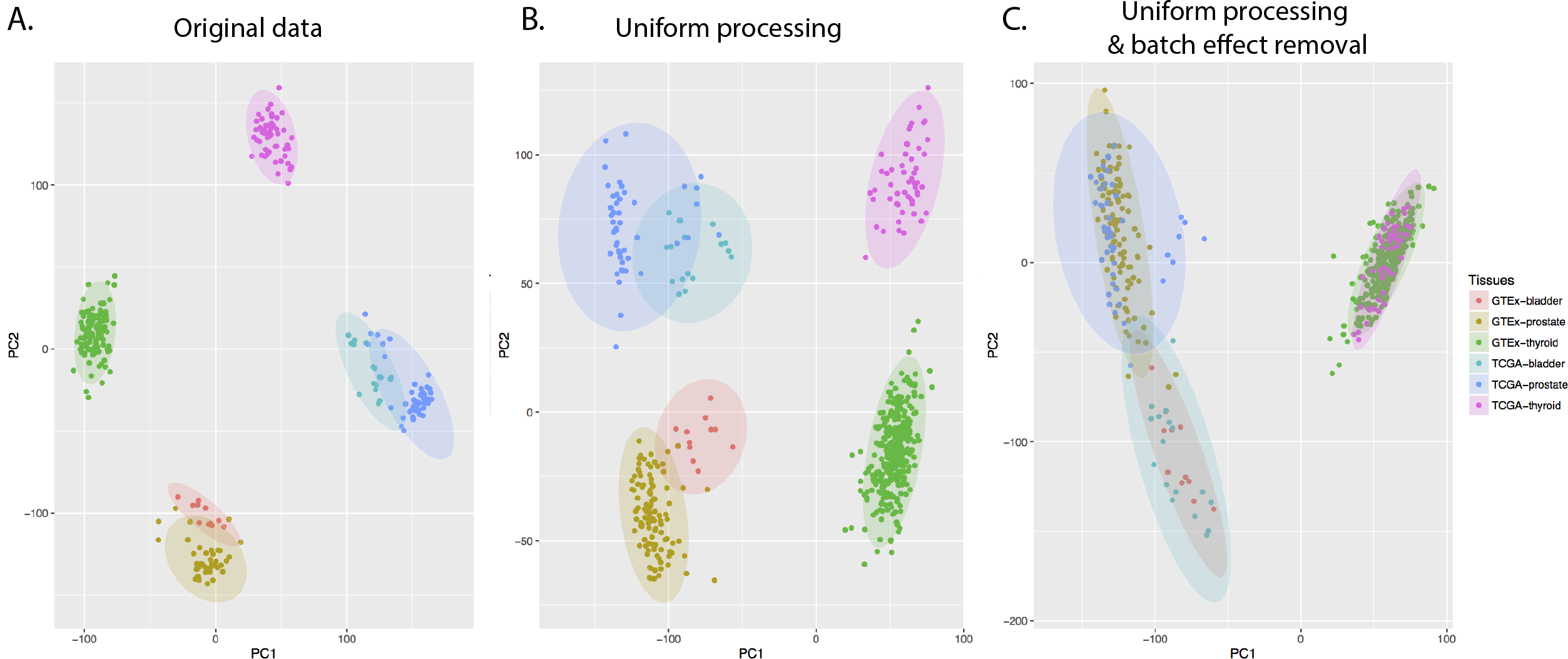
Effect of uniform processing and batch effect removal on expression levels in GTEx and TCGA. Two-dimensional plots are shown of principal components calculated by performing PCA of the gene expression values of bladder, prostate, and thyroid samples from GTEx and TCGA. **(a)** PCA of the level 3 data, i.e. the expression data from GTEx and TCGA. GTEx expression data was quantile normalized (see **Fig. S1B**). **(b)** PCA of the expression data after uniform processing through our pipeline, before batch bias correction. **(c)** PCA of the expression data after uniform processing through our pipeline, after batch bias correction.

However, uniform realignment and expression quantification using our pipeline did not fully resolve these differences; while the first principal component was now the tissue, the second principal component was still defined by the source (**Fig. 2B**), indicating that study-specific biases still accounted for significant variation in RNA-seq expression levels within each tissue type. This result shows that consistent realignment and expression quantification alone are not sufficient, and that further study-specific batch effects need to be removed in order to be able to compare expression data from TCGA and GTEx.

To this end, we next added a batch-effect correction step to our pipeline, using ComBat [12] (see **Methods**), which successfully corrected our example data and resulted in clustering by tissue type (**Fig. 2C**).

To determine whether uniform alignment and expression quantification was an essential step, or whether batch effect removal via ComBat by itself was sufficient, we also applied ComBat directly to the level 3 data from GTEx and TCGA (GTEx-quantified data was rescaled using quantile normalization). We found that batch effect removal by itself is not sufficient, and that the combination of uniform processing of sequencing reads followed by additional batch effect removal is required to make data from the TCGA and GTEx projects comparable (**Fig. S2**). We validated the expression similarities observed in the principal component analysis through hierarchical clustering (**Fig. 3**).

**Figure 3.**
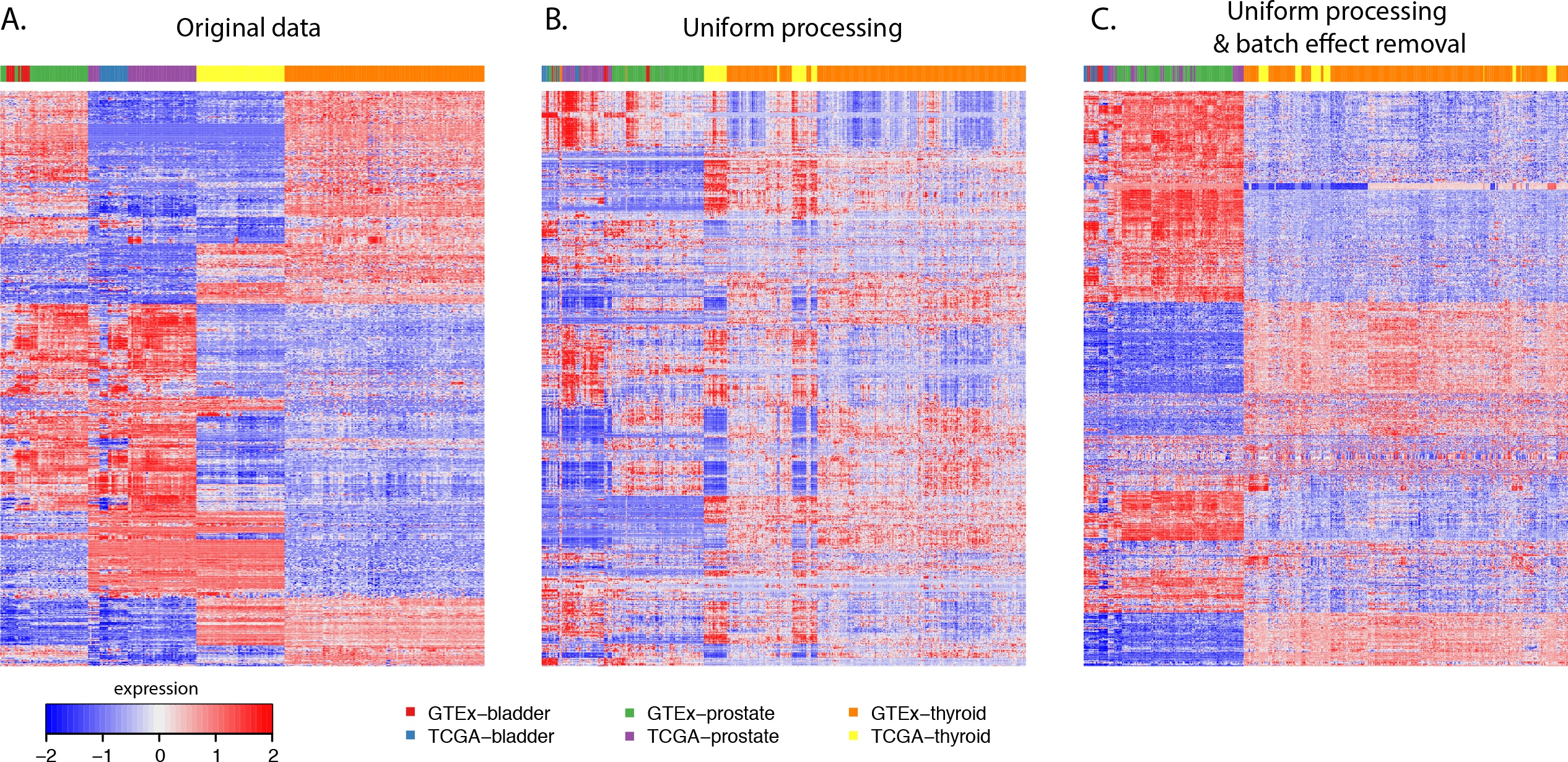
Hierarchical clustering of GTEx and TCGA bladder, prostate, and thyroid data shows the effect of uniform processing and batch effect correction. **(a)** level 3 expression data from GTEx and TCGA; **(b)** gene expression calculated using our pipeline prior to batch bias correction; **(c)** our expression data after batch bias correction.

Our results demonstrate that uniform realignment and expression quantification, together with explicit correction for study-specific biases, are not only effective, but also necessary for removing batch effects and making samples from different studies comparable.

Finally, we examined the expression levels of three cancer driver genes, ERBB2, IGF2, and TP53, in our batch-effect corrected data (**Fig. 4**). ERBB2 expression was significantly higher in a subset of tumor samples, consistent with the frequent amplifications observed in various tumor types. IGF2 showed a similar pattern, with a subset of tumor samples expressing the gene at levels several orders of magnitude higher than those in normal samples. TP53, on the other hand, is often affected by truncating mutations in cancer, which leads to decreased levels of RNA due to nonsense-mediated decay, an effect that is visible in the normalized RNA data.

**Figure 4.**
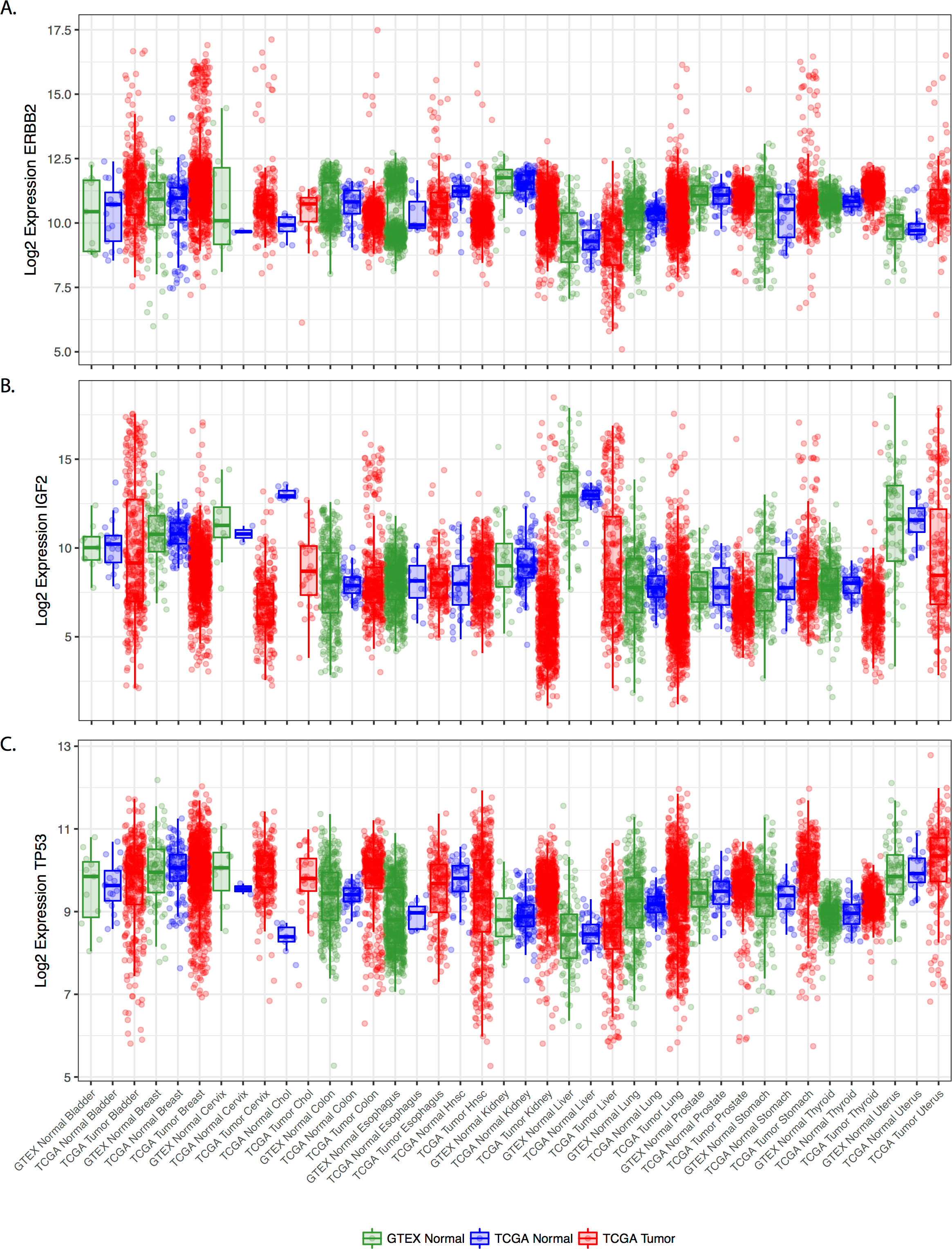
Normalized expression across tissue and cancer types for three known cancer genes: **(a)** ERBB2; **(b)** IGF2; **(c)** TP53.

The data generated using our pipeline has been deposited into GitHub at https://github.com/mskcc/RNAseqDB.

## Discussion

Recent large-scale studies, such as TCGA and GTEx, have resulted in an unprecedented volume of RNA-seq data. While these data offer new opportunities to identify the mechanisms underlying disease, the comparison of data from different sources poses a great challenge, due to batch effects inherent in the data from these studies.

Here, we describe a pipeline for correcting and integrating RNA-seq data across studies. Our pipeline starts with raw sequencing reads, performs uniform alignment and read quantification, and then removes study-specific batch effects. We have applied our pipeline to two of the largest studies in the field: GTEx and TCGA, processing 2790 normal RNA-seq samples from GTEx as well as 7576 samples from TCGA. Our results show that our pipeline is able to correct the biases specific to GTEx and TCGA, and, thus, make the samples from the two projects comparable.

Further efforts will be required to process samples for which there were not sufficient normal samples in the TCGA project, as well as GTEX samples for which there is no corresponding tumor type in the TCGA project. Our approach currently relies on the presence of normal samples in both studies that need to be integrated.

The data created has been deposited to GitHub and will be made accessible through the cBioPortal for Cancer Genomics [13], [14], so that investigators can conveniently mine the data and conduct integrative analyses. The resulting resource will benefit the research of cancer and other human diseases, as the pipeline can be used for the integration of RNA-seq data from other sources.

## Methods

### RNA-seq data

Raw paired-end reads of the RNA-seq samples for the TCGA project were retrieved from the Cancer Genomics Hub (CGHub, https://cghub.ucsc.edu). When FASTQ files were not available, e.g. for stomach adenocarcinoma, we downloaded aligned sequence reads (in BAM format) and extracted reads from BAM files with the Java program ubu.jar (https://github.com/mozack/ubu) before processing samples using our pipeline. GTEx samples were downloaded from the Database of Genotypes and Phenotypes (dbGaP, http://www.ncbi.nlm.nih.gov/gap), which hosts >9,000 RNA-seq samples (in SRA format) for the GTEx study.

### Analysis pipeline

We employed STAR aligner [15], a fast accurate alignment software used widely in the NGS community, to map reads to UCSC human reference genome hg19 and reference transcriptome GENCODE (v19), using recommended parameters, e.g. ‘–outFilterType BySJout’ and ‘–outFilterMultimapNmax 20’, etc., which are also standard options of the ENCODE project for long RNA-seq pipeline. Samples with alignment rates less than 40% were excluded from further analysis. The detailed parameters we used to run STAR and the codes of our pipeline are available at GitHub (https://github.com/mskcc/RNAseqDB).

The software tools FastQC, Picard (http://picard.sourceforge.net/index.shtml), RseQC [16], and mRIN [17] were used to evaluate sample quality. RNA degradation, as detected by mRNA, was present in some GTEx and TCGA samples. Since degradation can bias expression level measurements and cause data misinterpretation, we decided to exclude samples with evidence for degradation. To determine an appropriate degradation cutoff for mRIN, we used prostate cancer samples from the TCGA project, which had undergone extensive pathological, analytical, and quality control review and which had been shown to include a significant portion of degraded samples [11]. **Fig. S3** compares mRIN scores with RNA Integrity Numbers (RIN) calculated by TCGA for prostate samples [11]. It shows that mRIN is negatively correlated with RIN (Pearson correlation<-0.93). We used −0.11 as the degradation threshold (horizontal line in **Fig. S3**) for mRIN, which corresponds roughly to the cutoff 7.0 used by TCGA for RIN. Samples with mRIN<-0.11 were regarded as degraded and, thus, excluded from further analysis.

To verify mRIN’s performance on other tissues, we manually examined coverage uniformity over gene bodies for other tissues using the tool RseQC [16] and compared it with mRIN scores. We calculated the number of reads covering each nucleotide position and the average coverage for all long genes (>4000nt). **Fig. S4** shows the average coverage for TCGA prostate and bladder samples, each curve representing gene body coverage of a sample. In **Fig. S4A**, the 4 samples with the most uneven coverage are the ones deemed degraded in **Fig. S3**. We made similar observations in the other tissues examined, e.g. bladder in **Fig. S4B**, where the samples with the most imbalanced gene body coverage were the ones with the lowest mRIN scores. These results confirmed that mRIN is capable of measuring degradation for other tissues.

When running STAR, we specified an option ‘–quantMode TranscriptomeSAM’ to make STAR output a file, Aligned.toTranscriptome.out.bam, which contains alignments translated into transcript coordinates. This file was then used with RSEM [18] to quantify gene expression. The program “rsem-calculate-expression” in the RSEM package requires strand specificity of the RNA-seq sample, which is estimated using RseQC [16].

We also used another transcript quantification tool FeatureCounts[19] to generate integer-based read counts. Overall, the output from FeatureCounts was highly consistent with that of RSEM (Spearman correlation > 0.95). However, for genes with multi-mapping reads (i.e., reads mapped to multiple genes), FeatureCounts differs from RSEM and tends to underestimate expression levels in comparison with RSEM (because it discards multi-mapping reads). For example, the transcript from the PGA3 gene, which encodes human pepsinogen A enzyme that is highly abundant in the stomach, is identical to the transcripts of two other genes, PGA4 and PGA5. Its measurement in stomach by FeatureCounts (in default settings) is generally lower than that by RSEM (see **Fig. S5**). In our analysis below, we primarily used results by RSEM.

To ensure that TCGA normal samples remain comparable with TCGA tumors after removing batch biases from the normals, we also included TCGA tumors in our sample-gene matrix, which were processed in the same way as the normals using our pipeline from raw sequencing reads. In **Table S2**, we used bladder and lung as examples to show the parameters we used to run ComBat. As indicated in **Table S2**, we treated all TCGA samples, both tumors and normals (of the same tissue type) as one batch. To prevent ComBat from suppressing tumor-specific signals, we created a model to specify each sample to be either ‘normal’ or ‘tumor’, with which to run ComBat (see configuration file at https://github.com/mskcc/RNAseqDB/blob/master/configuration/tissue-conf.txt).

### Principal component analysis

To perform principal components analysis, we firstly remove genes with invariant expression levels and then log_2_-transform sample-gene matrix. Next, we utilize an R function ‘prcomp’ (with the ‘center’ option set to TRUE) to do principal components analysis. The two-dimensional PCA plot is created using an R function ‘autoplot’.

### Hierarchical clustering

For hierarchical clustering of expression data, we used the R function Heatmap.3 using default parameters (e.g., distance: euclidean, hierarchical clustering method: ward, etc.) as well as the top 1000 most variable genes in the data matrix.

---

**Figure S1. (a)** Ranges of GTEx and TCGA RNA-seq gene expression levels in bladder normal samples. **(b)** Gene expression levels in GTEx samples were scaled using quantile normalization.

**Figure S2.** PCA plot after applying quantile normalization and ComBat to the level 3 data of the 3 tissues, bladder, prostate, and thyroid, from GTEx and TCGA

**Figure S3.** Comparing mRIN with RNA Integrity Number (RIN) calculated by TCG for prostate cancer samples. The horizontal line at −0.11 is our cutoff. A sample with mRIN < −0.11 is deemed degraded. TCGA prostate cancer group used 7.0 (vertical line) as cutoff.

**Figure S4.** Gene body coverage of the TCGA prostate and bladder samples. Each curve in the figure represents average coverage of genes (from 5’ to 3’) in a sample. To ease visual examination, only long genes (>4000nt) were used in the calculation of the coverage and only the normal samples were plotted.

**Figure S5.** Expression of gene PGA3 in six tissues. Gene expression in (a) and (b) were quantified using FeatureCounts and RSEM, respectively. The same set of GTEx and TCGA (both tumor and normal) samples was used to compare FeatureCounts and RSEM for each tissue type.

**Table S1.**
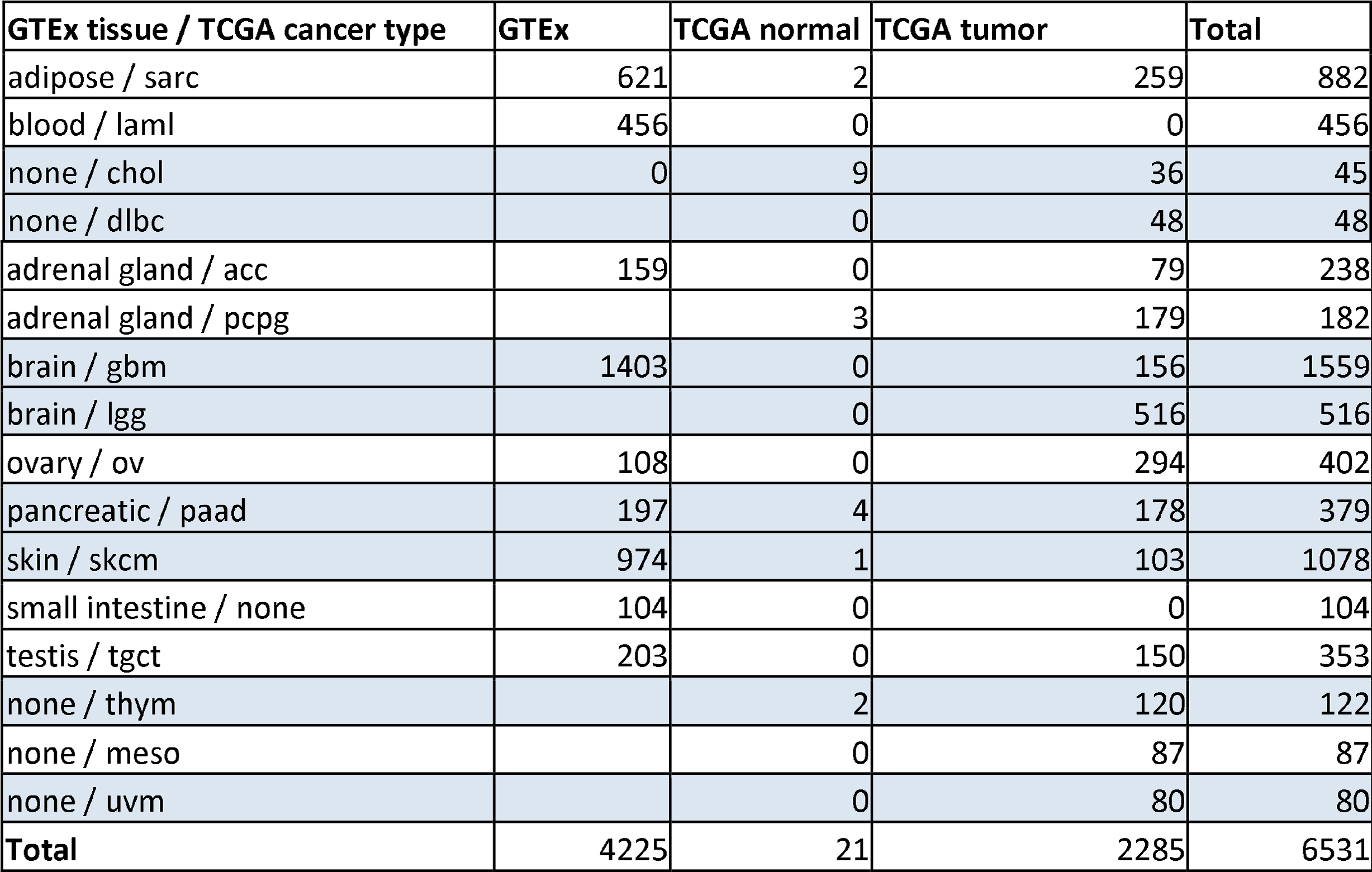
Samples with no or insufficient numbers of normal samples available in TCGA or GTEx.

**Table S2.** ComBat parameters for: (a) bladder, (b) lung. TCGA lung cancer has two subtypes: lung adenocarcinoma (LUAD) and lung squamous cell carcinoma (LUSC). We designated LUSC in same batch as LUAD. (a) Parameters of ComBat for bladder (b) ComBat parameters for lung.

|  | GTEX bladder | TCGA BLCA normal | TCGA BLCA tumor |
| --- | --- | --- | --- |
| **Batch** | 1 | 2 | 2 |
| **Variable of interest** | normal | normal | tumor |

**(b)** ComBat parameters for lung
|  | GTEX lung | TCGA LUAD normal | TCGA LUAD tumor | TCGA LUSC normal | TCGA LUSC tumor |
| --- | --- | --- | --- | --- | --- |
| **Batch** | 1 | 2 | 2 | 2 | 2 |
| **Variable of interest** | normal | normal | tumor | normal | tumor |

